# *zetadiv*: an R package for computing compositional change across multiple sites, assemblages or cases

**DOI:** 10.1101/324897

**Authors:** Guillaume Latombe, Melodie A. McGeoch, David A. Nipperess, Cang Hui

**Affiliations:** School of Biological Sciences, Monash University, Melbourne 3800, Australia; Department of Biological Sciences, Macquarie University, North Ryde, NSW 2109, Australia; Centre for Invasion Biology, Department of Mathematical Sciences, Stellenbosch University, Matieland 7602, South Africa; African Institute for Mathematical Sciences, Cape Town 7945, South Africa

**Keywords:** species turnover, alpha diversity, beta diversity, zeta diversity, occurrence data

## Abstract

Spatial variation in compositional diversity, or species turnover, is necessary for capturing the components of heterogeneity that constitute biodiversity. However, no incidence-based metric of pairwise species turnover can calculate all components of diversity partitioning. Zeta (ζ) diversity, the mean number of species shared by any given number of sites or assemblages, captures all diversity components produced by assemblage partitioning. *zetadiv* is an R package for analysing and measuring compositional change for occurrence data using zeta diversity. Four types of analyses are performed on bird composition data in Australia: (i) decline in zeta diversity; (ii) distance decay; (iii) multi-site generalised dissimilarity modelling; and (iv) hierarchical scaling. Some analyses, such as the zeta decline, are specific to zeta diversity, whereas others, such as distance decay, are commonly applied to beta diversity, and have been adapted using zeta diversity to differentiate the contribution of common and rare species to compositional change.

**Highlights:**

- An R package to analyse compositional change using zeta diversity is presented.
- Zeta diversity is the mean number of species shared by any number of assemblages
- Zeta diversity captures all diversity components produced by assemblage partitioning
- Analyses relate zeta diversity to space, environment and spatial scale
- Analyses differentiate the contribution of rare and common species to biodiversity

## 1. Introduction

### 1.1. Species turnover in practice

Spatial variation in compositional diversity, or species turnover, is one of the key properties for quantifying the components of heterogeneity that constitute biodiversity, along with total richness and measures of uniqueness, such as endemism and phylogenetic distinctiveness (Magurran and McGill, 2011). Species turnover can show a wide range of responses to environmental changes, and good conservation practice requires the understanding derived from its effective measurement and description for both species that are common and rare (McGeoch and Latombe, 2016; Socolar et al., 2016).

Despite the role of compositional dissimilarity (or similarity) in understanding biodiversity, no single measure previously connected the range of assemblage patterns constructed from species presence-absence data (Hui and McGeoch, 2014). Species turnover is traditionally measured by beta diversity, which quantifies compositional change between pairs of individual assemblages (Chao et al., 2012; Jost, 2007). To compare three or more assemblages, the mean of the pairwise similarities is often used (Jost et al., 2011). However, such incidence-based metrics of pairwise compositional change emphasize the differences in rare species composition between assemblages, and do not capture the characteristics of community structures caused by common species shared by many assemblages. Although multiple-site metrics have also been developed to quantify the heterogeneity in assemblage composition (Baselga, 2013; Diserud and Ødegaard, 2007; Ricotta and Pavoine, 2015), these measures rely on averaging non-independent pairwise values and are difficult to interpret.

### 1.2. Necessity of zeta diversity

Zeta (ζ) diversity, the mean number of species shared by any given number of sites or assemblages, was proposed as a metric to capture all diversity components produced by assemblage partitioning (Hui and McGeoch, 2014). Computing zeta diversity for combinations of sites from 2 to *n* sites (the orders of zeta), where *n* is the total number of sites, along with ζ_1_, the average number of species per site (i.e. alpha diversity), is necessary to obtain a mathematically comprehensive description of species assemblages, and cannot be achieved by only considering alpha and beta diversity. Let us consider a simple example with three sites containing 22 species each (i.e. ζ_1_ or α). Let us assume that each site shares exactly 10 species with any of the other two sites (i.e. ζ_2_ or β) (Figure 1). There are then multiple ways to partition species diversity between the three sites. At one extreme, there may be no species shared by the three sites simultaneously (ζ_3_ = 0). At the other extreme, the 10 species shared by any two sites may actually be extremely common and be shared by all three sites (ζ_3_ = 10) (Figure 1). These different partitions of species diversity therefore correspond to very different species assemblages, whereas they have the same alpha and beta diversity values. Many other examples are possible, where alternative diversity partitions exist even with the same alpha, beta and gamma diversity values.

**Figure 1.**
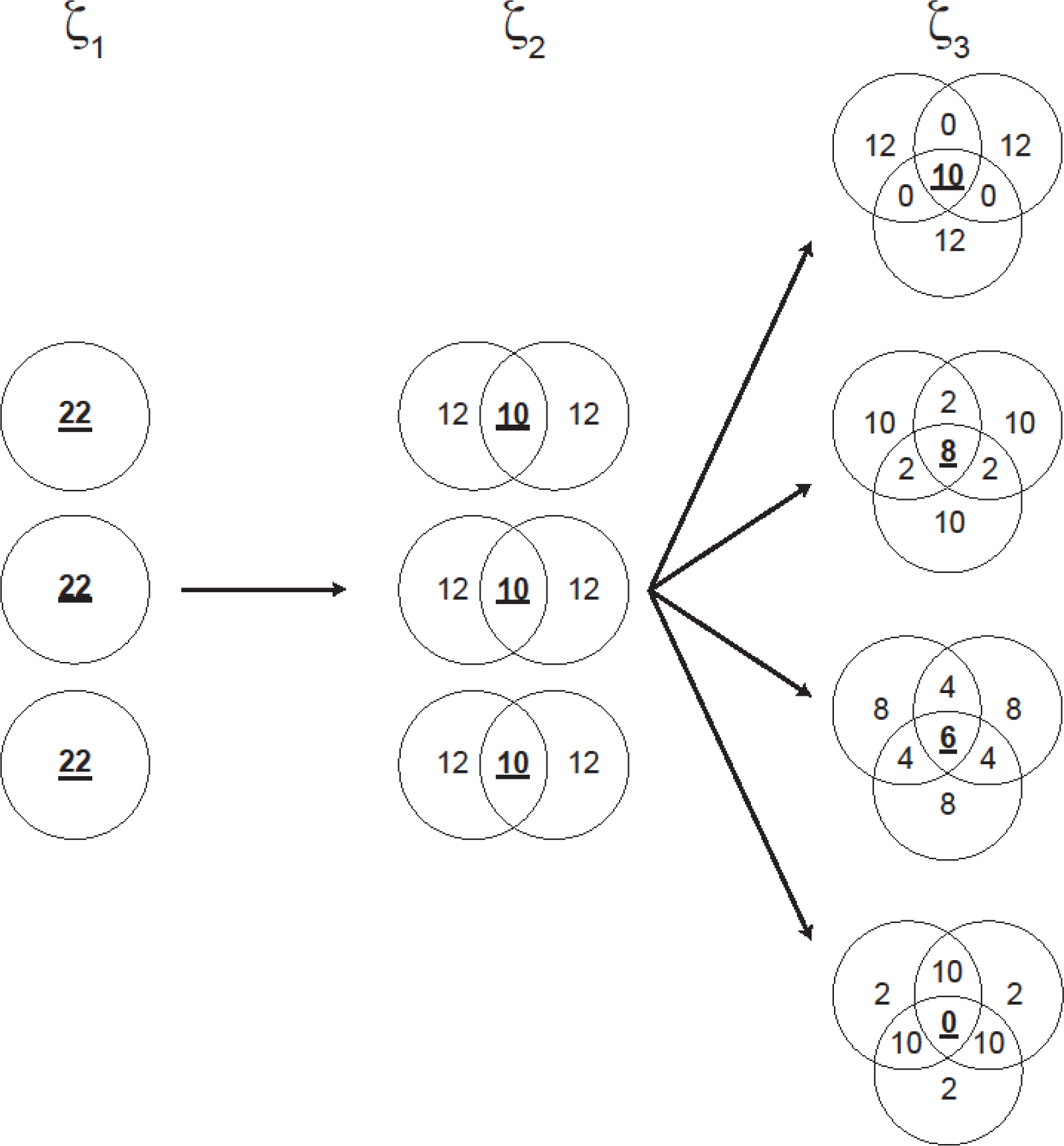
Four (amongst many) different ways to partition species turnover when 3 assemblages (zeta order = 3) are combined. Only considering richness (alpha diversity, corresponding to ζ_1_) and pairwise compositional change (beta diversity, corresponding to ζ_2_, where zeta order = 2) provides an incomplete description of the community. Numbers in bold and underlined are the values of zeta diversity for orders 1 to 3.

As a consequence of the comprehensive description provided by zeta diversity, as outlined by Hui and McGeoch (2014), zeta diversity enables the computation of a broad range of existing diversity metrics, and the quantification of continuous change in biodiversity over landscapes. For example, species accumulation curves, endemic-effort relationships and occupancy-frequency distributions can all be derived from zeta diversity. Importantly, from examining an extensive dataset of 291 communities, Hui and McGeoch (2014) identified two most common parametric forms of zeta diversity decline with the increase in the number of given sites – negative exponential and power law, which together account for 80% of examined communities and may differentiate stochastic from deterministic assembly processes, respectively (see for example Roura-Pascual et al., 2016). By providing a common currency for measuring biodiversity from occurrence data, zeta diversity provides an avenue for understanding the mechanistic basis of spatial patterns in diversity. This includes examining if environmental change affects rare and common species differently, or testing hypotheses about the relative importance of deterministic versus stochastic assembly processes in generating patterns of biodiversity.

Because it links different community patterns together, zeta diversity can be used for identifying community assembly processes. The identification of processes generating community assemblages usually relies on community patterns (e.g. Dornelas et al., 2006; Latombe et al., 2015). Since multiple assembly processes can generate the same community pattern, multiple patterns are needed to provide a more comprehensive description of the community and discriminate between processes (Grimm et al., 2005; Grimm and Railsback, 2012). Using multiple, different patterns can nonetheless generate bias due to possible redundancy between their information content (Latombe et al., 2011). Since multiple incidence-based patterns can be derived from zeta diversity, zeta diversity offers a powerful basis for discriminating between community assembly processes while avoiding issues of pattern redundancy. Following this logic, zeta diversity has been used to compare and provide insights on the nature of compositional change over space and time using 10 datasets encompassing a whole range of levels of biological organisation at various spatial and temporal scales, including birds, insects, plants, microbes, and crop pests, but also intracellular processes in humans, showing the potential of zeta diversity for describing and unveiling the functioning of systems beyond classical site-by-species structure (McGeoch et al., 2017).

Importantly, zeta diversity enables the contribution of rare and common species to compositional change to be disentangled. On average, common (widespread) species are more likely to be present in any site and to be shared by any two sites than rare species. The variation in the number of species shared by different pairs of sites are therefore mostly driven by rare species, and so are analyses based only on alpha and beta diversity. By contrast, since rare species cannot, by definition, be shared by many sites, differences in zeta values for high orders of zeta is only driven by common species. Although conservation actions are mostly orientated towards rare species, common species are getting more attention (McGeoch and Latombe, 2016) as their importance for ecosystem functions is increasingly recognised (Gaston, 2010). Understanding the contribution of common and rare species to species turnover is therefore necessary. In practice, defining the distinction between rare and common species is subjective and must be done for each species individually. By contrast, zeta diversity calculates the contribution of species from rare to common as a continuum, avoiding multiple and largely subjective decisions.

### 1.3. Aims and novelty of the *zetadiv* R package

Here we introduce the *zetadiv* package for R (R CoreTeam, 2013). The *zetadiv* package (available on CRAN; https://CRAN.R-project.org/package=zetadiv) was created to measure and analyse compositional change for occurrence data using zeta diversity. The functions of the *zetadiv* package can be categorised into four kinds of analyses described in detail in the following (Appendix A, Table A1): (i) the analysis of zeta diversity decline explores how the number of species shared by multiple assemblages decreases with increasing number of assemblages within combinations, and what information is contained in the form of this decline; (ii) the analysis of the distance decay of zeta diversity illustrates how zeta diversity for different orders varies with distance between sites; (iii) Multi-Site Generalised Dissimilarity Modelling (MS-GDM, an adaptation of Generalised Dissimilarity Modelling; Ferrier et al., 2007), computes the contribution of different environmental variables and distance to zeta diversity for different orders; (iv) the analysis of the hierarchical scaling of zeta diversity unravels how zeta diversity varies with grain.

Analysis of the decline in zeta diversity uses an incremental increase in the numbers of assemblages included in the combinations. It is therefore an application unique to zeta diversity as it combines alpha and beta (ζ_1_ and ζ_2_) with higher orders of zeta in a single analysis, to provide a comprehensive description of species turnover. By contrast, as we detail below, the other three kinds of analyses have in the past been applied using beta diversity, and other R packages exist to compute such analyses. The *vegan* package (Oksanen et al., 2018) enables the computation of a wide range of beta diversity measures and of the hierarchical scaling of beta diversity with sampling grain. The *simba* package (Jurasinski and Retzer, 2012) enables the comparison of different slopes of the distance decay of beta similarity. Generalised Dissimilarity Modelling can be performed on beta diversity using the *gdm* package (Manion et al., n.d.). As we illustrate in the examples below, the *zetadiv* package extends such analyses and enables their application to zeta diversity for selected numbers of assemblages beyond pairwise beta diversity (n ≥ 2; see the full R code in Appendix B for fully reproducible examples and figures).

## 2. Biodiversity data

All the functions of *zetadiv* require at most four types of data (except for the functions that use the outputs of other functions, such as plotting functions). (i) Occurrence data, in the form of sites-by-species (rows-by-columns) data frames, are required by all functions. (ii) When spatial information is needed, a data frame with the projected or geographical coordinates of the sites or assemblages can be used. (iii) Instead of the spatial coordinates, a distance matrix between sites, independently computed, can be provided, when measures of connectivity other than Euclidian or orthodromic (i.e. distance between two points on the globe defined by their geographic coordinates) distance are required (e.g. Manhattan distance or distance accounting for the path of least resistance). (iv) A site-by-variable data frame representing the environmental variables of the sites or assemblages can be provided for MS-GDM analysis.

Two datasets, describing two different ecosystems, and complying with these requirements are included in the package to demonstrate the functions (Appendix A, Table A2). The first dataset is an inventory of resident, terrestrial bird survey data (presence-only) from the BirdLife Australia Atlas of Australian Birds (1998-2013) and covering South-East Australia (Barrett et al., 2003). The species occurrences are complemented by maps of environmental variables for the same region, including proportion of natural environments, irrigated agriculture and plantations, as well as human density, water features (www.abs.gov.au), temperature and precipitation (www.worldclim.org; Fick and Hijmans, 2017), and elevation (www.gebco.net). The bird and environmental data were arranged into two continuous grids at two different spatial scales (same spatial extent but a fine grain [25 × 25 km grid cells] and a coarse grain [100 × 100 km grid cells], therefore producing two scales. Only cells whose richness was within 10% of estimated asymptotic richness are included in the datasets (Latombe et al., 2017), to limit the occurrence of false absence. The second dataset is an inventory of the presence and absence of springtails and mite species in 12 plots (4 transects and 3 altitudes) on Marion Island, along with the altitude of the sites and the side of the island where they are located (McGeoch et al., 2008; Nyakatya and McGeoch, 2008).

In the following, we use the fine-grain bird data to illustrate the four different kinds of analyses. The fine-grain data, together with the seven environmental variables, can be loaded using the following commands:

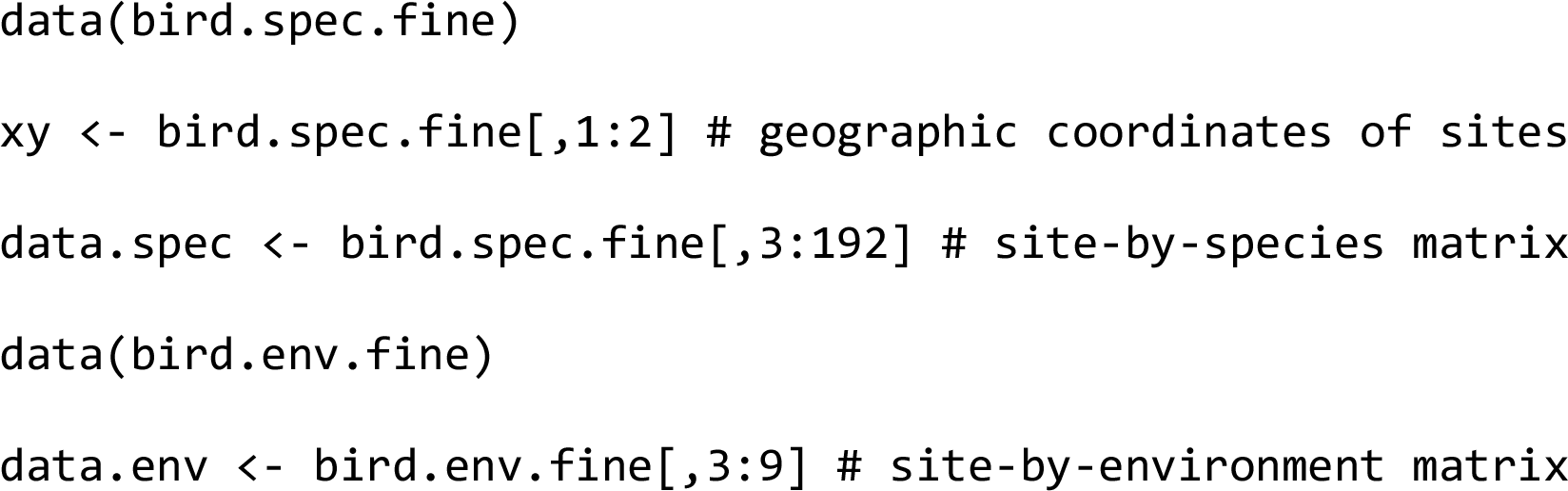

## 3. Zeta diversity and zeta decline

### 3.1. Description

The functions `Zeta.order.ex` and `Zeta.order.mc compute` ζ_i_, the number of species shared by any *i* assemblages (the order of zeta) in two alternative ways. `Zeta.order.ex` computes the expected value of zeta diversity for order *i*. Let *P_j_* be the probability of species *j* with occupancy *O_j_* occurring in *i* given sites out of the *N* surveyed sites. The expected value can be calculated as the sum of the probability over all species *S*:

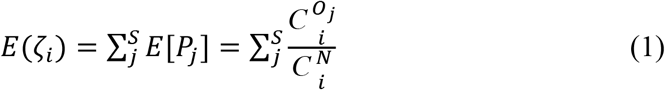

where 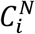 and 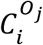 are binomial coefficients giving the total number of possible combinations of *i* sites out of a total of *N* or *O_j_*, respectively. The variance is then given by the summation of the covariance of the probability:

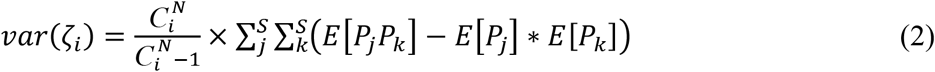

where

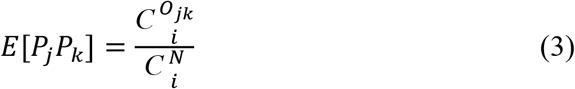

and *O_jk_* is the number of sites in which both species *j* and *k* are present (also referred to as joint occupancy; Hui, 2009). The number *O_ij_* corresponds to the element *ij* of the S×S dimensional matrix **M**^T^**M**, where **M** is the site(row)-by-species(column) matrix of occurrence and T matrix transposition. Note that the variance in Equation 2 is corrected for bias using Bessel’s correction (Kenney and Keeping, 1951), which corresponds to the default in `Zeta.order.ex`. This is suitable if the assemblages represent a sample of the total study system. In case of a continuous grid sample or in lab experiments, for which the incidence data can be exhaustive, the exact variance can also be computed by setting `sd.correct = FALSE` in the function parameters.

By contrast, `Zeta.order.mc` (for “Monte Carlo sampling”) computes zeta diversity by averaging the number of shared species for *i* assemblages over all possible combinations of the *i* assemblages from *N* total assemblages. The shared species for *i* assemblages is obtained using the dot product of species (1/0) vectors. When all possible combinations are used, `Zeta.order.mc` and `Zeta.order.ex` are equivalent. For large *N* and intermediate *i*, the number of combinations for *i* assemblages, 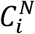, becomes very high, and the computational complexity becomes intractable. The user must therefore provide a value `sam`, representing the number of samples over which ζ_i_ should be computed. If 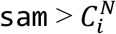, ζ_*i*_ is computed exactly, but otherwise approximated over `sam` random combinations. The impact of sam on the computation of ζ_i_ can be assessed using the function `Zeta.sam.sensitivity` (Appendix A, Figure A1).

In contrary to `Zeta.order.ex`, for each combination *j* of *i* assemblages, `Zeta.order.mc` allows for the computation of a normalised version zeta (i.e. *ζ_ij_*/*S_j_*), where *S_j_* is either (i) the total number of species over the assemblages in the specific combination *j* (i.e. the gamma diversity of the combination *j*, therefore equivalent to the Jaccard similarity index), (ii) the average number of species per assemblage in the specific combination *j* (i.e. the alpha diversity of the combination *j*, therefore equivalent to the Sørensen similarity index), or (iii) the minimum number of species over the assemblages in the specific combination *j* (therefore equivalent to the Simpson similarity index). Normalised zeta may be suitable when richness varies widely across regions or systems being compared.

The formulas described above for `Zeta.order.ex` and `Zeta.order.mc` correspond to combinations of any *i* assemblages over all assemblages. This subsampling scheme may nonetheless not be the most appropriate for some data. For example, the turnover of assemblages arranged in a linear fashion along a gradient (e.g. Rivadeneira et al., 2002; Whittaker, 1956) may be better analysed by combining assemblages close to each other, and using a specific assemblage as a reference (Whittaker, 1967). Several sub-sampling schemes are possible in `Zeta.order.mc`. Assemblages can be combined using a nearest-neighbour approach to explore patterns of local turnover. When a nearest-neighbour approach is used, the combinations can be non-directional, or directional, moving away from a fixed-point origin or a fixed-edge origin (for example for ecological systems being invaded from a specific direction) (McGeoch et al., 2017). A focal assemblage plus the closest (*i*-1) assemblages are then be used for calculating ζ_*i*_. The focal assemblage can be the fixed-point origin or any other assemblage. There are therefore 4 possible subsampling schemes, whose pertinence depends on the specific study (see McGeoch et al., 2017 for additional details and a comparison of the zeta declines using different sub-sampling schemes for the well-known Smokey Mountain dataset of Whittaker 1956, 1967): the ALL scheme using combinations of any assemblages (the default scheme), the non-directional nearest neighbour (NON) scheme, in which each site is associated to its *i*-1 nearest neighbours to compute ζ_*i*_, the directional nearest neighbour using a specific assemblage or an edge as a reference (DIR), and each site is associated to its *i*-1 nearest neighbours in the opposite direction to the reference to compute ζ_*i*_, and the fixed-point origin (FPO) scheme, in which a specific assemblage is always combined with its *i*-1 nearest neighbours to compute ζ_*i*_ (i.e. similar to NON but using one specific assemblage only). When the FPO is located outside of the study area, it corresponds to a fixed-edge origin (FEO) scheme, in which assemblages close to the edge are combined with their nearest neighbours.

The functions `Zeta.decline.ex` and `Zeta.decline.mc` then compute the values of ζ_*i*_ for a range of orders *i* (Figure 2; Appendix A, Figure A2). As the number of assemblages increases, the number of shared species amongst assemblages necessarily decreases, hence a decline in zeta. These functions also compute the ratio ζ_i_/ ζ_i-1_, which is called the retention rate and quantifies the proportion of species that are retained in additional samples. The retention rate is especially useful to reveal features of the zeta decline that are indistinguishable from the observation of the decline itself, allowing for highlighting differences in the structure of compositional change between datasets or study areas, and for detecting spatial structure in gradients of vegetation when using different sub-sampling schemes (McGeoch et al., 2017).

**Figure 2.**
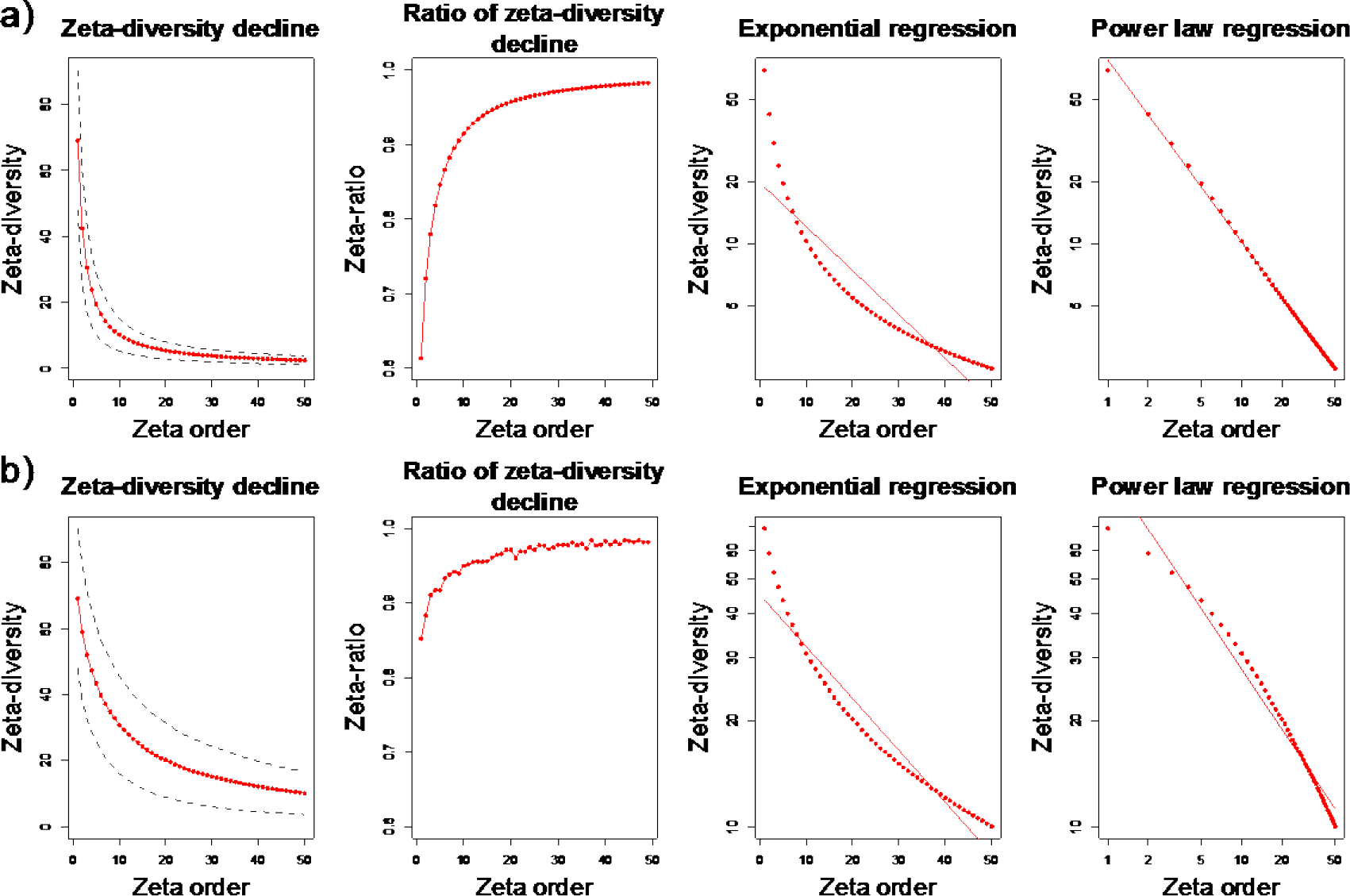
Zeta decline between orders 1 to 50 characterising the bird data for 25 × 25 km cells computed with a) `Zeta.decline.ex` (‘expected’, i.e. all combinations) and b) `Zeta.decline.mc` with `sam=1000` and using a non-directional nearest-neighbour (NON) subsampling scheme.

Finally, an exponential and a power law parametric form are fitted to the zeta decline. These are the two most common parametric forms observed in nature (Hui and McGeoch, 2014). The parametric form of the decline may signal the relative roles of stochastic or deterministic assembly processes, although it may also be affected by assemblage richness and sample size. The function `Plot.zeta` plots the outputs of `Zeta.decline.ex` and `Zeta.decline.mc`.

Computing zeta diversity for different orders has been used, for example to validate the outputs of self-organising maps used for pest profile analyses, which group together areas with similar profiles of species composition (Roigé et al., 2017). Pairwise comparisons of sites enables the identification of clusters with few shared species and therefore high uncertainty. Using orders of zeta beyond pairwise comparisons enables to further refine the uncertainty level of the remaining clusters by distinguishing between clusters with low (i.e. superficial) and high similarity for higher orders of zeta.

### 3.2. Example

The zeta decline of bird species over South-East Australia was computed from orders 1 to 50 using the ALL and the NON subsampling schemes across grid cells and plotted using the following commands (the seed is set to 1 for reproducibility):

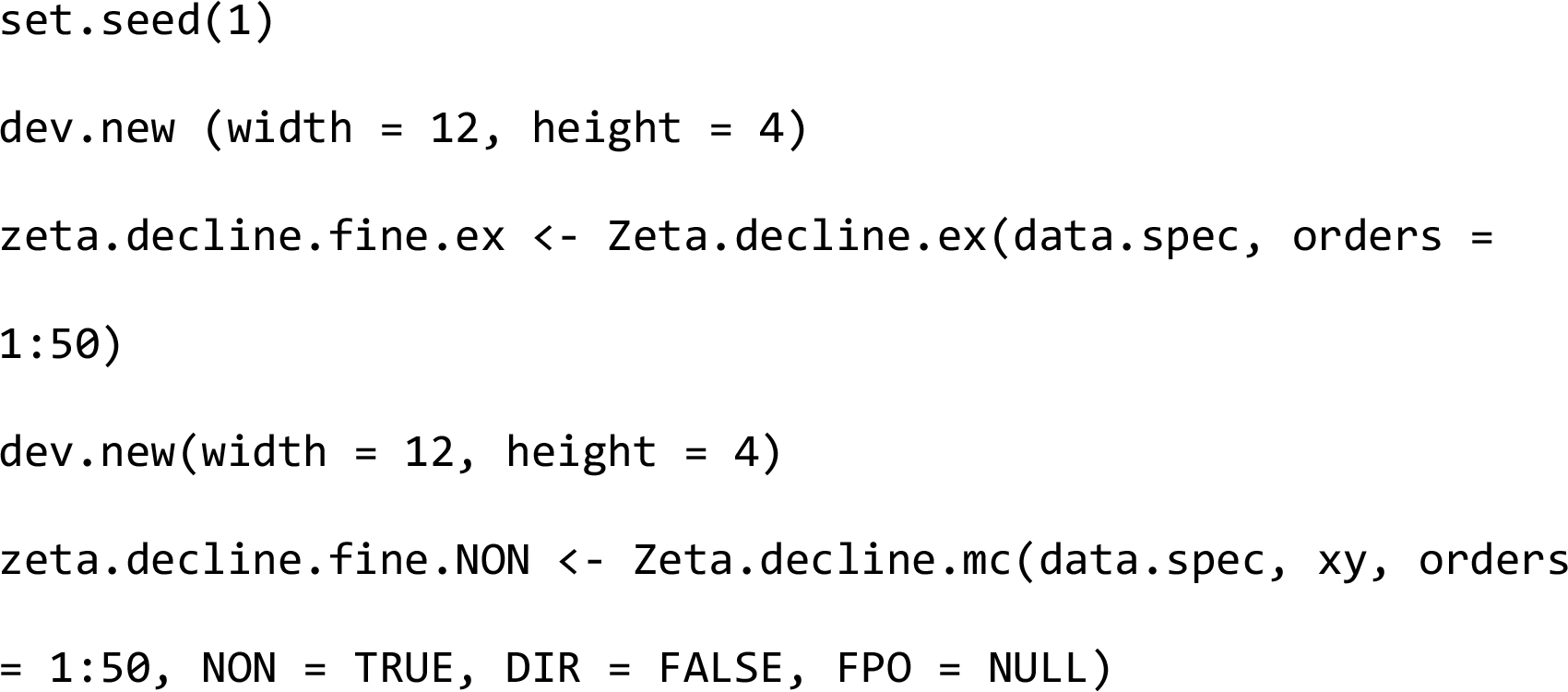

The `NON = TRUE` parameter indicates that the NON scheme must be used. If the `FPO` parameter contains coordinates, they take precedence over the `NON` parameter. If `DIR = FALSE`, the FPO (or FEO) scheme applies. The DIR scheme requires both `DIR = TRUE` and a set of coordinates in `FPO`.

Comparing outputs of the ALL and the NON subsampling schemes provides information on the effect of the spatial scale on species turnover. When all cells are combined, the zeta decline better fits a power law than an exponential parametric form (Figure 2a; ΔAIC = 270.61), therefore suggesting that species are distributed in a deterministic fashion across South-East Australia. The retention rate (ζ_*i*_ / ζ_*i*-1_) increases steadily, but starts levelling off after 20 assemblages, indicating that, below that value, few species are retained as new assemblages are considered, but many more are, proportionally, beyond 20 assemblages. The asymptote therefore provides an indication of the scale at which species can be considered to be rare and common.

When the cells are combined using the NON scheme, the retention rate is higher than for the ALL scheme for low orders of zeta, indicating that the zeta values decline at a lower rate. This suggests some level of spatial aggregation of species, with closer cells sharing more rare species (and common species to a lesser extent), as can be expected. The zeta decline computed with the NON scheme is also better fitted by a power law rather than exponential parametric form.

## 4. Distance decay of zeta

### 4.1. Description

The distance decay of similarity is a well-known community descriptor (Morlon et al., 2008; Nekola and White, 1999), i.e. as distance between assemblages increases, two assemblages are expected to become less similar and to share fewer species. Typical research questions that can be addressed by considering the distance decay of zeta-diversity include: (i) the explicit distances over which species assemblages differ; (ii) how do the decay patterns of rare and common species differ, providing insight on the spatial properties of their distributions.

The function `Zeta.ddecay` generalizes distance decay and enables its computation for any number of assemblages. For many sites, it uses the same Monte Carlo sampling as `Zeta.order.mc`, and can therefore be applied to normalised zeta. For more than two assemblages, distances between assemblages (either computed from sites coordinates or from a custom distance matrix) must be combined for each combination of sites, for example as the mean distance across *n* sites. The function is flexible and enables users to define how they should be combined, using a built-in or a custom function (see Latombe et al., 2017 for a discussion on the impacts of using different functions). `Zeta.ddecay` regresses ζ_*i*_ over this measure of distance using three types of regression: (i) a generalized linear model, the default being linear regression, allowing constraints on the signs of the coefficients (ii) a generalized additive model (GAM), to allow for non-linearities and periodicities in the distance decay (Soininen et al., 2007) and (iii) a general additive model under shape constraint, or “shape-constrained additive model” (SCAM; Pya and Wood, 2015), set by default to a monotonically declining GAM. Additional options enable the definition of thresholds for distance which may be desirable, for example, for discarding uninformative long tails that would artificially make the slope of the distance decay in linear models more shallow. It is also possible to specify how to transform spatial distance according to any function. The function `Zeta.ddecays` calls `Zeta.ddecay` and computes the slope of the distance decay using linear models for different orders of zeta, and plots changes in slope as the order increases (Appendix A, Figure A3).

The distance decay of zeta can also be applied to time series of species composition, using time instead of distance, therefore computing a time decay of zeta diversity. Time decay of zeta diversity has been used to show differences in the response of bird communities of two different river basins to drought (McGeoch et al., 2017).

### 4.2. Example

The distance decays of ζ_2_, ζ_3_, ζ_5_, and ζ_10_ were assessed using a linear regression and a GAM (set with reg.type, whose default is linear regression) using the following commands (for `order = 2, order = 3, order = 5` and `order = 10`):

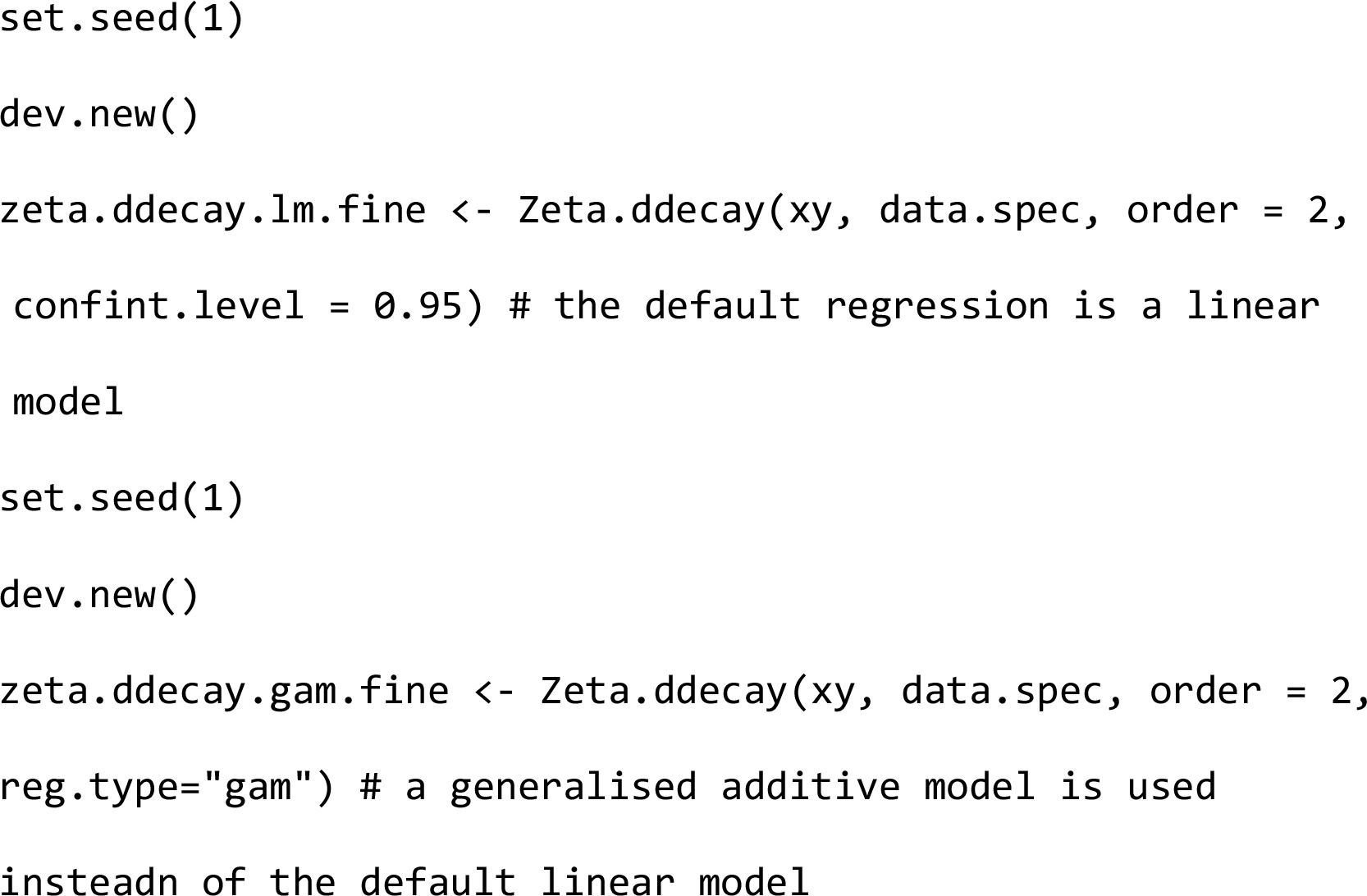

Both methods show a clear distance decay, even for ζ_10_, although it becomes less pronounced for high orders of zeta (Figure 3). The distance decay is more pronounced for ζ_3_ and ζ_5_ than for ζ_2_ (p-values = 0.001, 0.003; the significance was computed using the `diffslope2` function from the *simba* R package, Jurasinski 2012; see R code in Appendix B for details) suggesting that within the extent of this study, rare species are dispersed relative to the space-filling properties of the species with higher occurrence levels. The GAM shows that the distance decay is not linear for ζ_2_ and ζ_3_. In particular, ζ_3_ allows for the detection of a threshold at ~800 km after which the effect of distance on compositional change mostly disappears (Figure 3).

**Figure 3.**
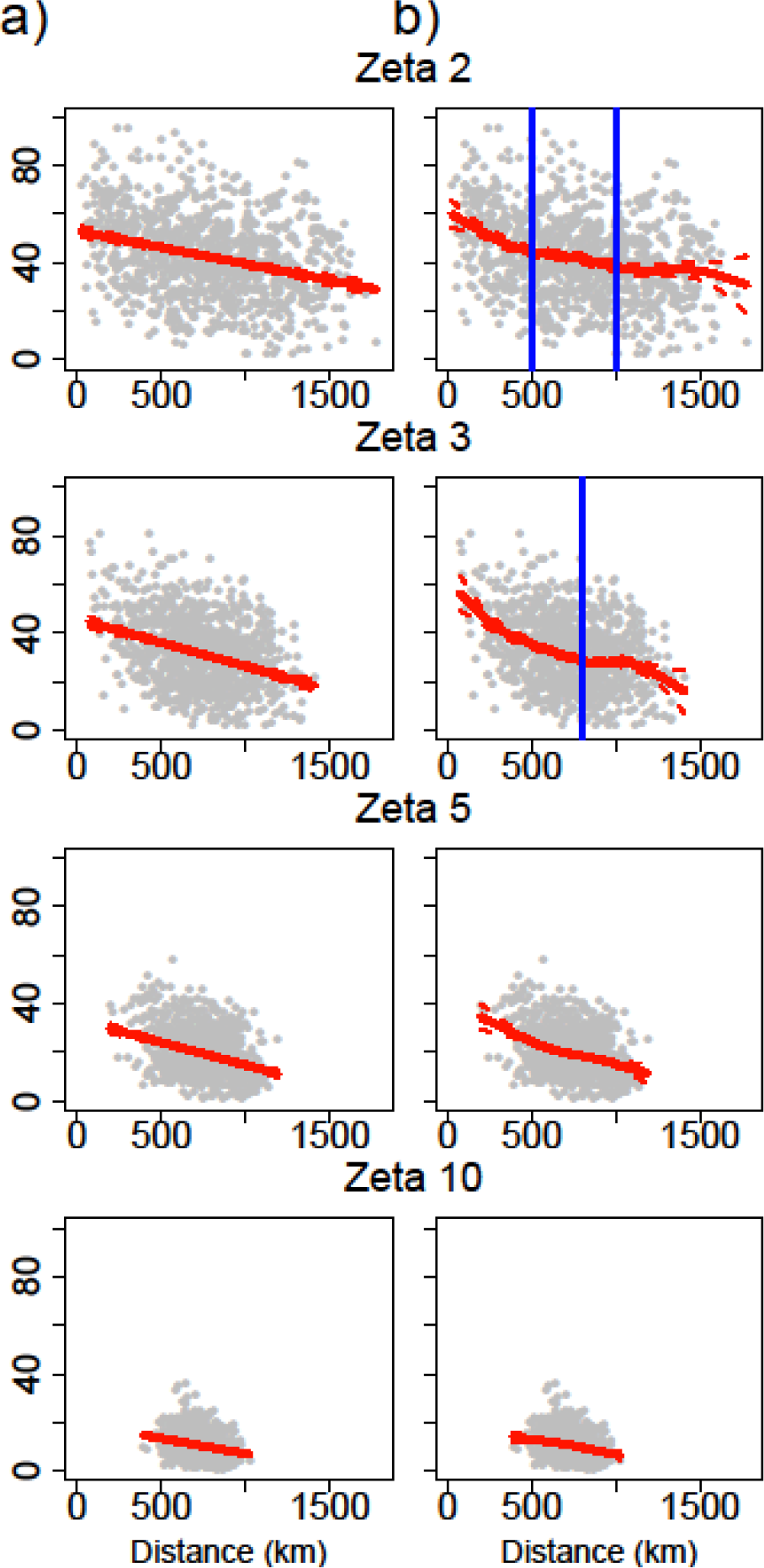
Distance decay of ζ_2_ to ζ_10_ for the bird data for 25 × 25 km cells using a) a linear regression and b) a generalised additive model (GAM). The linear regression shows clear distance decay, with a steeper slope for ζ_3_ and ζ_5_ than for ζ_2_, suggesting that rare species are dispersed relative to the space-filling properties of the species with higher occurrence levels. The GAM also reveals three slightly different rates of decline for zeta ζ2 (with thresholds at ~500 km and ~1000km) and two clearer different rates of decline for zeta ζ3 (with a threshold at ~800 km), indicated by the vertical blue lines.

## 5. Multi-site generalised dissimilarity modelling

### 5.1. Description

Multi-Site Generalised Dissimilarity Modelling (MS-GDM; Latombe et al., 2017) is inspired by Generalized Dissimilarity Modelling (GDM; Ferrier et al., 2007), a statistical technique for analysing and predicting changes in beta diversity from pairwise differences in environmental variables and spatial distance between sites using regression techniques. Following the same principles, the function `Zeta.msgdm` enables the regression of rescaled (ζ_*i*_/ζ_1_) or normalised ζ_*i*_ values (Jaccard, Sørensen or Simpson versions) over environmental differences and distance between assemblages. Since ζ_*i*_ is the number of species in common across *i* sites, we call it Multi-site Generalised Dissimilarity Modelling (see Latombe et al., 2017 for details). MS-GDM enables the inclusion of both continuous and categorical environmental variables as predictors. In the latter case, the environmental difference between *i* sites is computed as the number of different values across the *i* sites (and the maximum value is therefore *i*; Latombe et al. *in review*). MS-GDM also enables the inclusion of the zeta values of the same order from another group of species as predictors, when both groups are expected to be related to each other (such as native and alien species; Latombe et al. *in review*). Typical research questions that can be addressed by MS-GDM include: (i) whether variation in the number of shared species (compositional similarity) between assemblages is explained predominantly by either environmental differences or distance; (ii) whether the relative importance of different environmental variables and distance differs for rare and common species (by comparing low and high orders of zeta)

Four different types of regression techniques have been implemented: generalized linear models (GLM), with possible constraint on the sign of the coefficients, GAMs, SCAMs, and, following Ferrier et al. (2007), a combination of I-spline and GLM with constraints on the signs of the coefficients (see Latombe et al., 2017 for details). I-splines (Ramsay, 1988) are a kind of monotone spline functions that are used to transform the data before applying a generalized linear model with non-negative coefficients. This transformation accommodates non-linear relationships between zeta diversity and changes in environmental variables, but also the fact that the impact of change in an environmental variable may depend on the values of this variable (for example a change of temperature near the limit of the species thermal tolerance may have more impact on species occurrence than the same change in the middle of the range of its thermal tolerance).

The order of the I-splines and the number of knots (for the GAM, SCAM and I-plines) can be set by the user. The number of knots must be chosen carefully, as too many knots may result in overfitting (Manion, 2009). Moreover, as for any regression analysis, variables suffering from multicollinearity (e.g. VIF>10) should be removed (Dormann et al., 2013). As for the distance decay, for many sites, Zeta.msgdm uses the same Monte Carlo sampling as `Zeta.order.mc.` When *i* > 2, the environmental differences and distances between assemblages must also be combined for each combination, for example using the mean of differences.

A function `Ispline` to transform data using I-splines is also included in the package. Using the output from `Zeta.msgdm`, the function `Predict.msgdm` predicts the zeta values for new environmental data. The function `Plot.spline` is used to plot the I-splines for the different variables. Finally, the function `Zeta.varpart` computes variation partitioning (Legendre, 2008) for a model computed with `Zeta.msgdm`, to determine which part of ζ_*i*_ is explained by the environmental or the distance variables. `Zeta.varpart` uses the adjusted R^2^, to account for the use of several environmental variables, whereas distance is a single variable. Note that the non-adjusted R^2^ is computed as 1 − (residual sum of squares) / (total sum of squares), and makes sense only for linear regression, for which the residual sum of squares is normally distributed. Results of variation partitioning for the other regression techniques should therefore be interpreted with caution. In variation partitioning, some partitions may be negative (Legendre and Legendre, 2012). The function `Pie.neg` therefore considers negative values as 0 to plot the results as a pie diagram.

### 5.2. Example

Similar to Latombe et al. (2017), MS-GDM was computed for the Sørensen ζ_2_ and ζ_10_ using I-splines (and a binomial family with a log link, which requires a negative intercept, as shown by `cons.inter = −1`), as well as variation partitioning for linear regressions (contrary to MS-GDM, no constraint was applied on the sign of the regression by setting `method.glm = “glm.fit2”` so that the residuals are normally distributed, as explained above), using the following commands (for `order = 2` and `order = 10`):

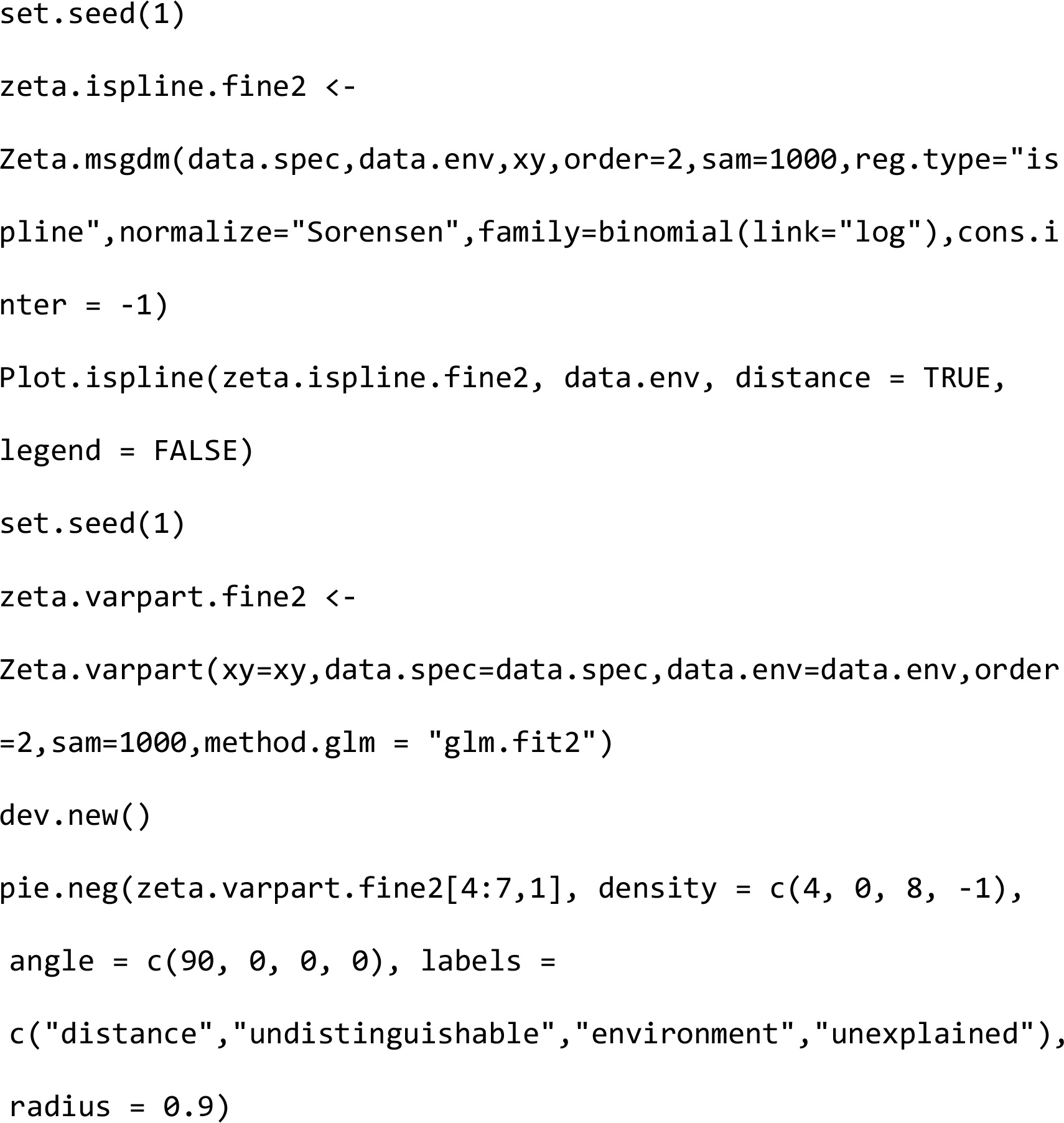

In these data, precipitation is the main predictor of bird compositional change for ζ_2_, especially for dry environments (as shown by the steep slope of the I-spline for low precipitations), followed by distance (Figure 4). For ζ_10_, which, contrary to ζ_2_, excludes the contribution of the rarest species to turnover, the importance of temperature and area per person increases. The decrease in the relative importance of precipitation may be due to the fact that common species are more likely to find refugia in areas containing water bodies during dry periods, whereas rare species may be more vulnerable to rainfall heterogeneity (discussed in further detail in Latombe et al., 2017). Results are slightly different from Latombe et al. (2017) because the Sorensen version of zeta was used here instead of just rescaling the zeta values by the overall ζ_1_, and different indices consider the influence of richness on turnover differently (Baselga, 2010).

**Figure 4.**
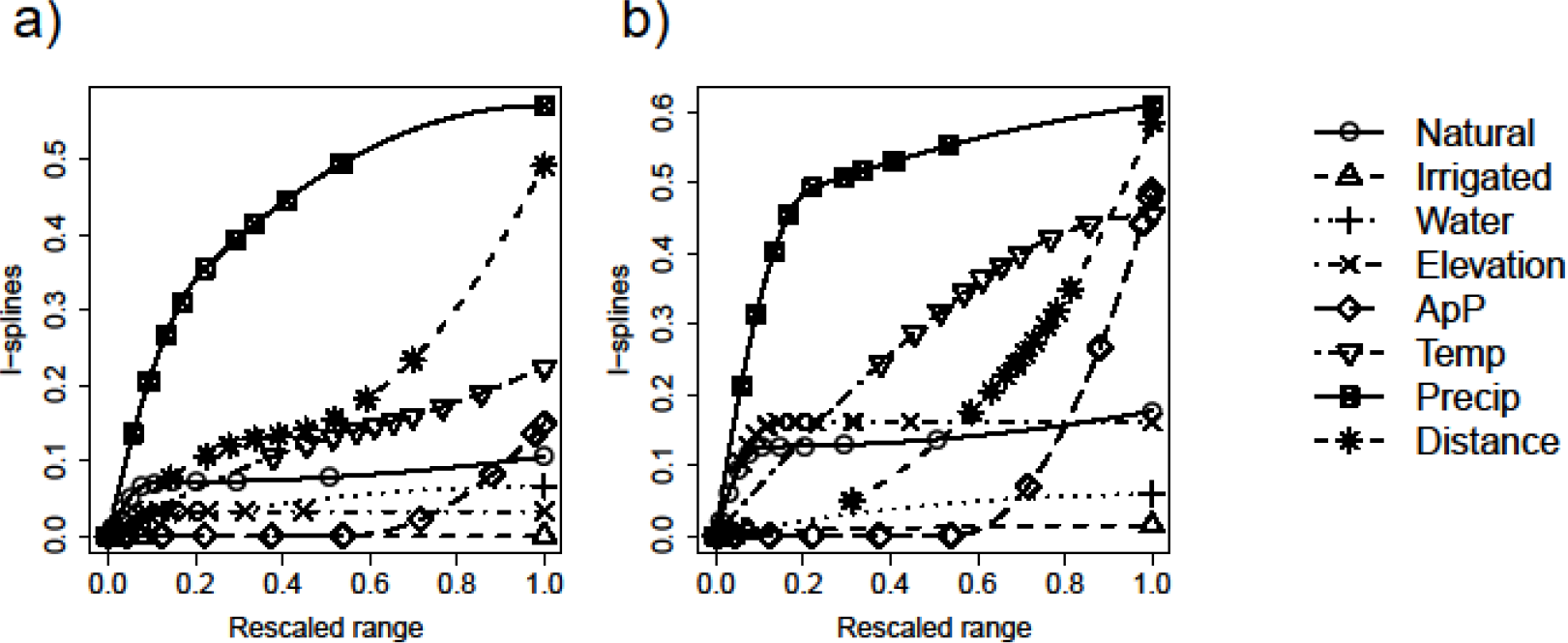
I-splines explaining zeta diversity of bird assemblages over South-East Australia for (a) ζ_2_ and (b) ζ_10_, using 7 environmental variables and spatial distance, for 25 × 25 km cells. The relative maximum values of the splines indicate the relative contribution of each variable to explaining zeta diversity. By contrast, the slope of the splines provide information on how the influence of each variable changes along the gradient of values. For example, changes in precipitation have more influence on compositional change in dry areas (low rescaled range value) than in wet areas (high rescaled range value), especially for ζ_10_ (Latombe et al. 2017).

Variation partitioning on ζ_2_ and ζ_10_ using I-splines and simple linear regressions shows that variation partitioning explains a larger proportion of variance for low orders of zeta than for high ones, indicating that the spatial distribution of rare species is more predictable than for common species (Figure 5). As expected, the I-splines explain a larger part of variations compared to linear regressions for both ζ_2_ and ζ_10_, due to their flexibility. In addition, these results linking environment with species compositional change rather than distance support the interpretation of the fact that the decline of zeta diversity is better fitted by a power law than by an exponential parametric form, suggesting deterministic community assembly.

**Figure 5.**
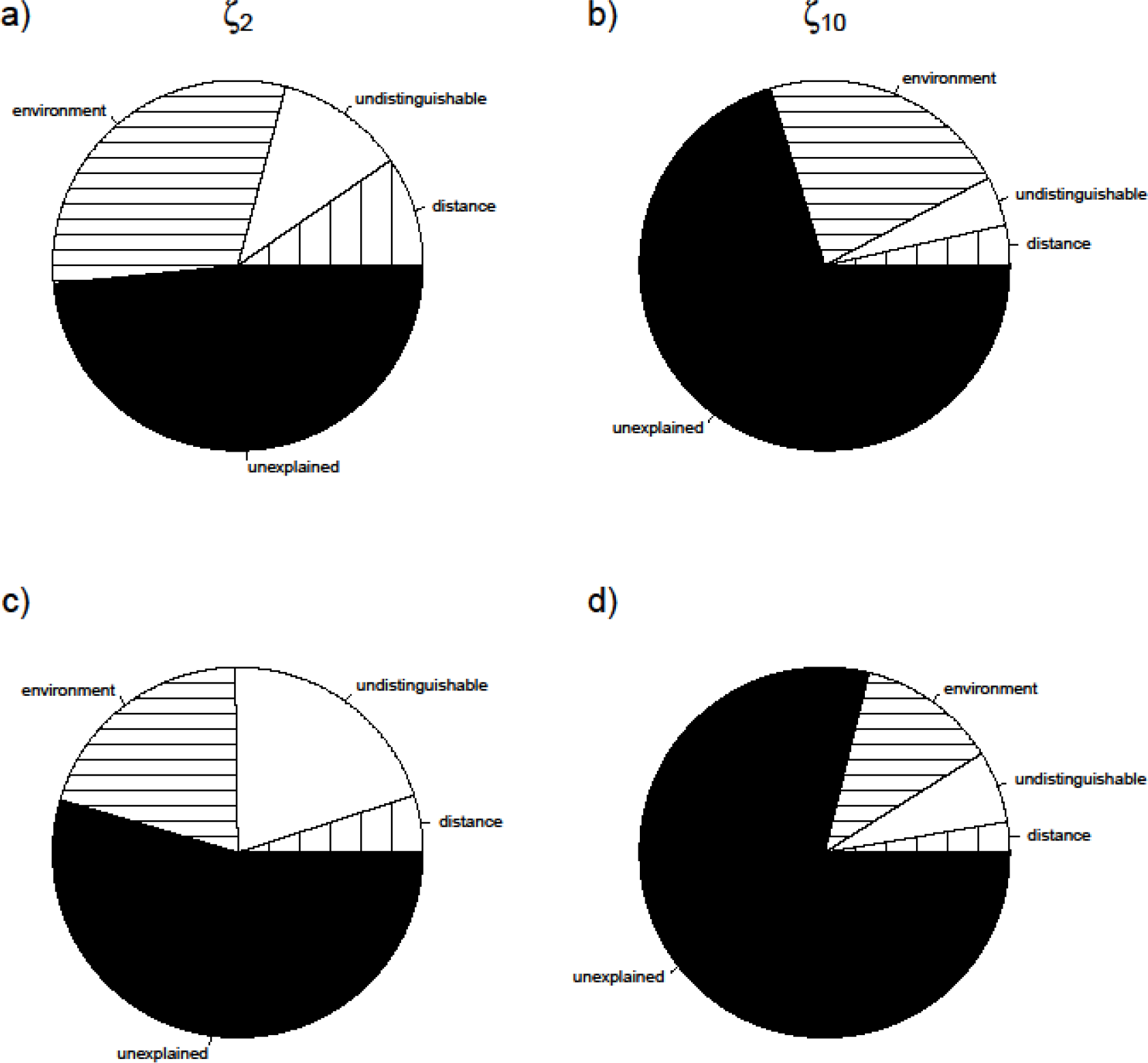
Proportion of variation (variation partitioning) in ζ_2_ and ζ_10_ explained by environmental and distance variables for the bird data for 25 × 25 km cells, when I-splines (a,b) and linear regressions (c,d) are used. The larger proportion of variation explained by the environment is consistent with the relative amplitudes of the corresponding I-splines (Figure 4).

## 6. Hierarchical scaling of zeta

### 6.1. Description

Like all biodiversity metrics, zeta diversity is sensitive to scale, *i.e.* to grain and extent (Hui and McGeoch, 2014). For compositional change, grain type follows three general sampling schemes (Scheiner et al., 2011): (i) sites arrayed as cells in a contiguous grid, (ii) sites arrayed as cells in a regular but non-contiguous grid and (iii) irregularly distributed sites of potentially varying size, such as islands (Figure 6). For data based on regular grids, the effect of scale can be assessed by grouping assemblages with their immediate neighbours (Figure 6a,b). For irregularly distributed areas, assemblages are grouped based on the distance between them (Figure 6c).

**Figure 6.**
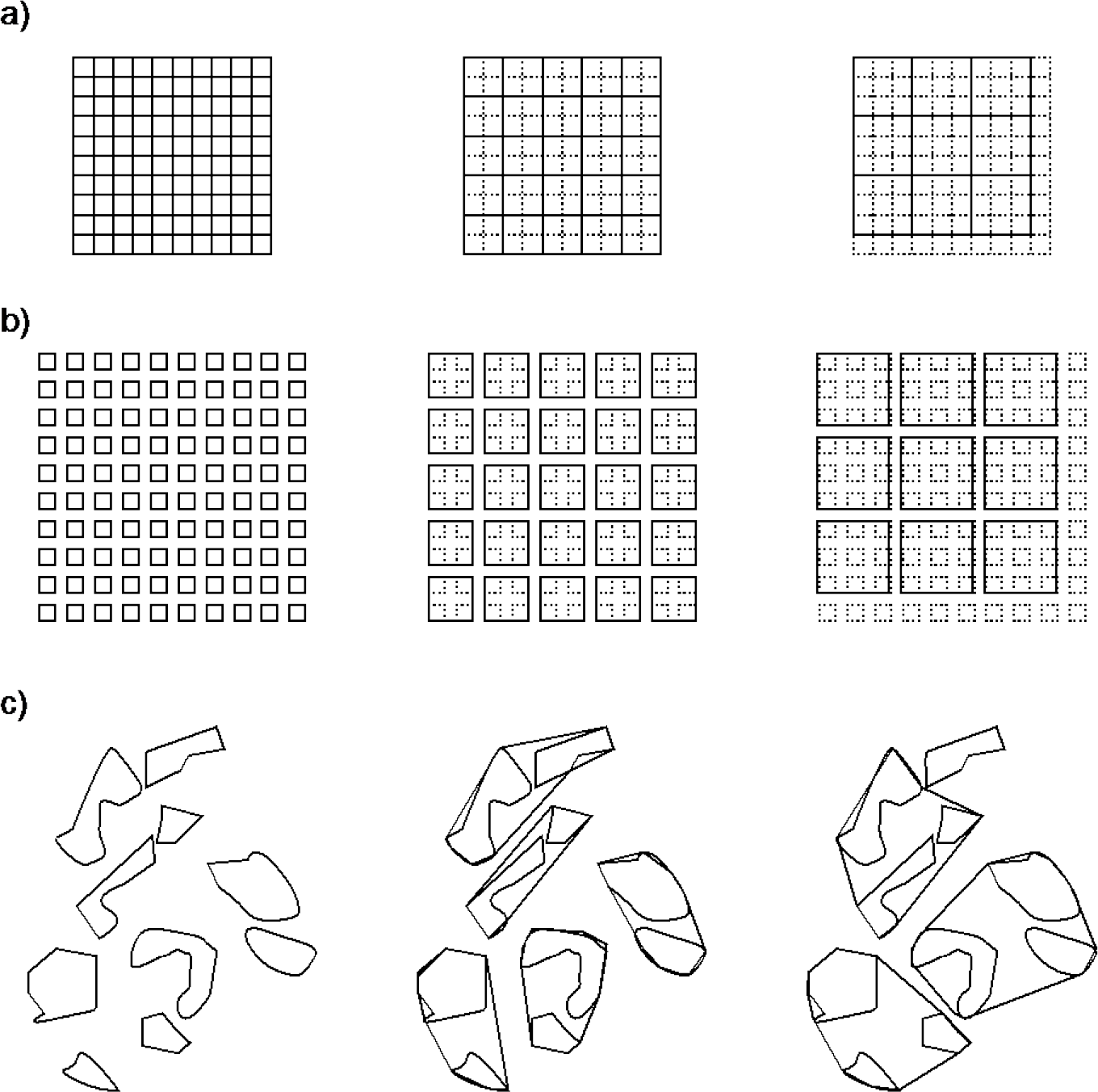
Resampling of data depending on the initial sampling scheme, classified according to three of the four sampling schemes defined by Scheiner et al. (2011) (the fourth type, strictly nested quadrats, is not relevant here) (see also McGeoch et al., 2017). For a) a continuous grid and b) a regular but discontinuous grid, adjacent cells are grouped together. For c) irregularly distributed sites of potentially varying size, such as islands, sites are grouped based on the distance between them. For a) and b), resampling based on minimum distance can also be applied to the grid cells, but grouping adjacent sites is not applicable to c). For a) and b), if the number of cells at fine grain is not a divisor of the number of cells at coarse grains, some cells are lost during aggregation. For c), the order in which sites are grouped can influence the final configuration. The bird datasets can be seen as a) and c), since the original cells are regularly distributed, but only cells with observed richness within 10% of estimated richness are included and the remaining cells are therefore irregularly distributed, but have a constant area.

When defining commonness based on the relative occupancy of a species (the number of sites or cells where the species is present divided by the total number of sites) (see McGeoch and Latombe, 2016), the proportion of rare species necessarily decreases as grain increases, whereas the proportion of common species increases. This is because the relative occupancy of a species necessarily increases (or stays constant) as grain increases. The rate at which rare species become more common with coarser grain depends on their spatial distribution (are species clustered or not) (Hui et al., 2010; Hui and A McGeoch, 2007; McGeoch and Gaston, 2002). For example, a species present in 4 adjacent cells arranged in a square (i.e. highest possible level of clustering) in a n × n continuous grid (Figure 6a) has an occupancy of 4/n^2^, and an occupancy of 1/(n/2)^2^ = 4/n^2^ once the grain is doubled if all four cells are combined into a single one. Any other spatial arrangement of the four cells will therefore generate an occupancy higher than 4/n^2^.

However, from a community perspective, species commonness and rarity are relative notions (McGeoch and Latombe, 2016). For a given occupancy, a species will be common in a community in which other species have a lower occupancy, and conversely. As we showed in the description of the zeta decline, the shape of the species retention rate across orders of zeta enables defining the threshold at which species can be considered to be rare or common. The distinction between rare and common species can therefore vary differently across scales for communities with different spatial arrangements of their species (McGeoch and Gaston, 2002). Species with different levels of spatial clustering will therefore contribute differently to the various orders of zeta diversity depending on the grain of the study. Species that are spatially dispersed will contribute to higher orders of zeta when the grain becomes coarser than species that are spatially clustered (Figure 7).

**Figure 7.**
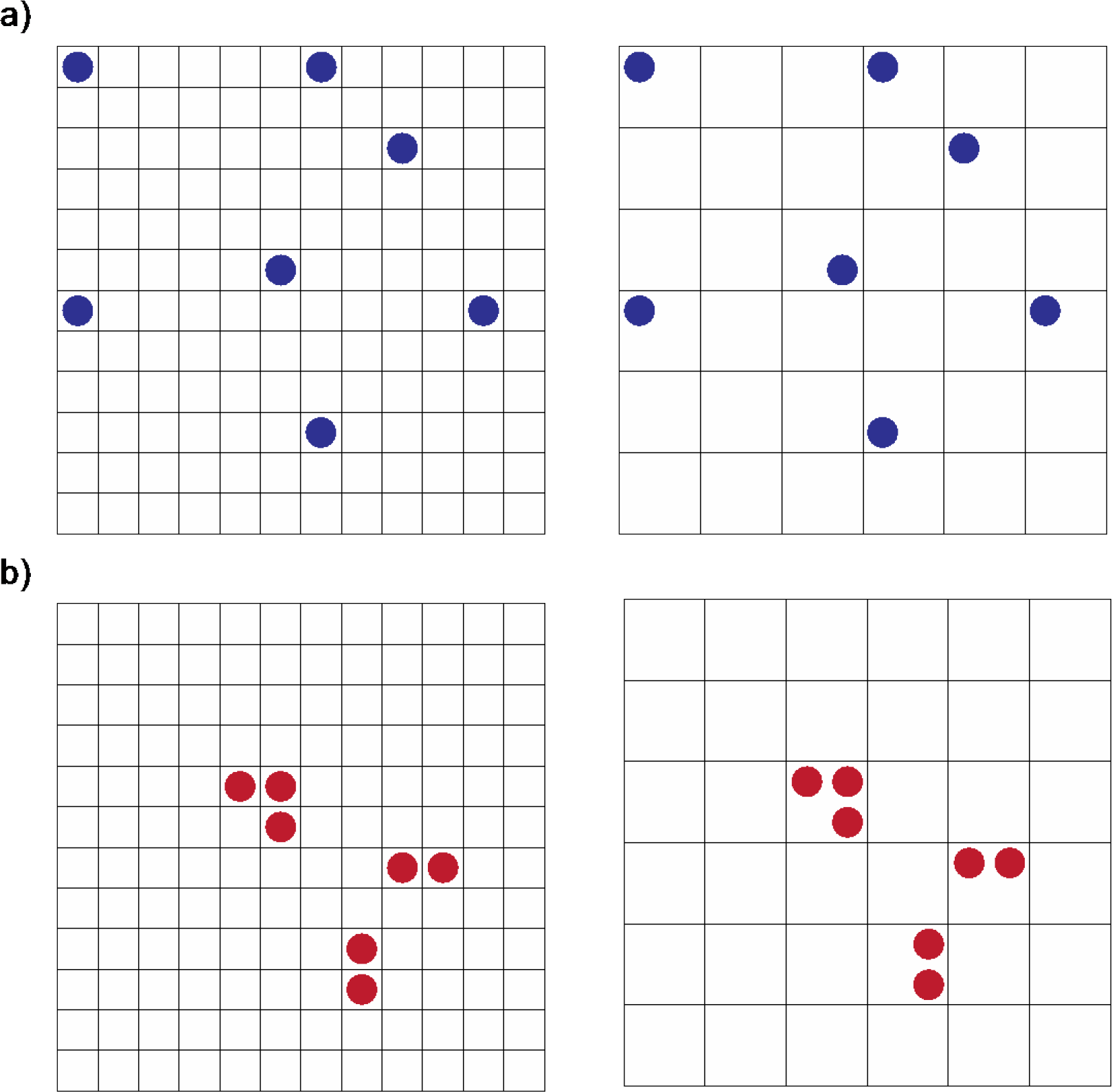
Effect of species spatial aggregation on their contribution to different orders of zeta when the grain changes. a) Highly dispersed species still contribute to high orders of zeta under the ALL sampling scheme (orders 1 to 7 in this example, since the species is present in 7 different grid cells) when the grain becomes coarser. b) At fine grain, spatially aggregated species contribute to higher orders of zeta (orders 1 to 7) than at coarse grain (orders 1 to 3).

Typical research questions that can be addressed by exploring the hierarchical scaling of zeta diversity therefore include: (i) how the characteristic of being common or rare varies with grain; and (ii) whether the sampling effort is sufficient to comprehensively study species turnover of both common and rare species.

In the *zetadiv* package, the functions `rescale.regular` and `rescale.min.dist` aggregate the species occurrence data, and combine the environmental data and the coordinates following a user-specified function such as the mean, based on the neighbours and on minimum distance, respectively, for a specific level of aggregation. The functions `Zeta.scale.regular` and `Zeta.scale.min.dist` compute ζ_*i*_ for a specific order *i*, for a range of levels of aggregation for the two methods. For `rescale.min.dist` and `Zeta.scale.min.dist`, the assemblages are aggregated iteratively: given a list of assemblages in a specific order, the first assemblage is combined with the closest ones, then the next available assemblage is combined with the closest available ones, and so on. Since the order of the assemblages in the list can impact the outcome of the algorithm, the function `Zeta.scale.min.dist` performs the analyses several times for each order and returns the mean.

### 6.2. Example

We assessed the hierarchical scaling of ζ_1_ to ζ_10_ by aggregating the 25 × 25 km cells (from 1 to 10 cells and then to 60 cells by steps of 10, as stated by `m = c(1:10,seq(20,60,10))`) based on minimum distance (Figure 6c) using the following commands (for order = 1 to order = 10):

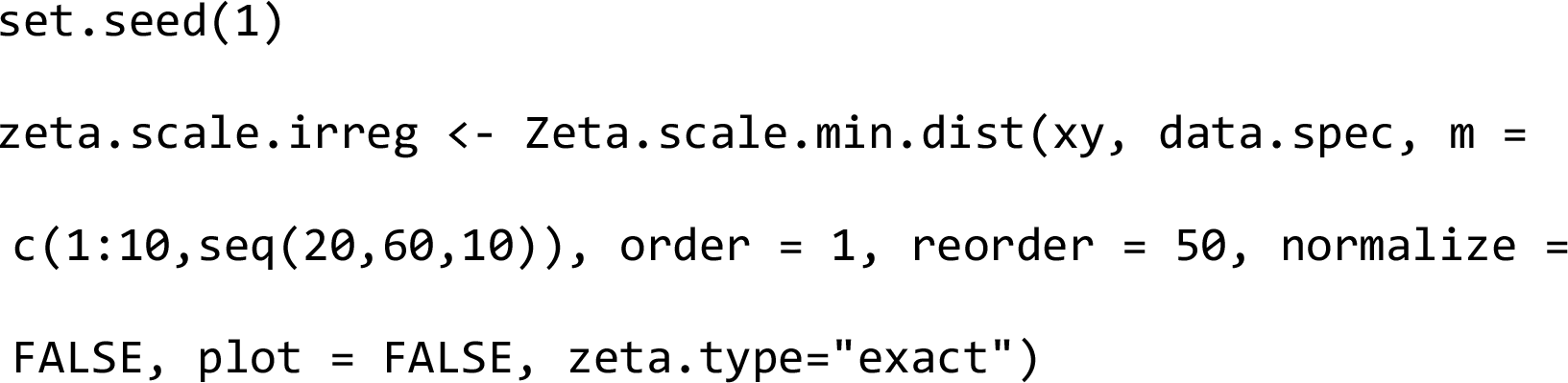

Since the order in which the cells are aggregated can change the results, the aggregation is performed 50 times (`reorder = 50`) and the average zeta values are computed.

As expected, zeta values increase as grain increases for all orders of zeta (Figure 8a). We also compared (ζ_i_- ζ_i-1_) for each grain, to compare the rates of increase across orders of zeta. Although zeta diversity increases with grain in a similar fashion for all orders (Figure 8a), the difference between the zeta values of different orders changes with grain and between orders. ζ_1_- ζ_2_ always decreases as the grain increases (Figure 8b), and the zeta decline becomes more shallow between orders 1 and 2 (Figure 8c). That is, less rare species are lost when increasing grain. By contrast, for higher orders of zeta, differences in the rates of increase between two consecutive orders of zeta increases when grouping 2 or 3 cells, then decreases (Figure 8b). This means that the zeta decline is steeper across orders 2 to 10 when aggregating 2 or 3 cells than for the fine grain data and for aggregating many cells (Figure 8c). These results suggest that a spatial grain of ~1250 km^2^ (~35 × 35 km) may be appropriate to study bird communities over Australia, as the sharper and more steady decline of zeta diversity indicates a more gradual distinction between common and rare species than at finer and coarser grains, for which the zeta decline becomes more shallow as the zeta order increases (Figure 8b,c). The ~1250 km^2^ grain may therefore be related to the scale at which bird species of different levels of rarity aggregate in South-East Australia.

**Figure 8.**
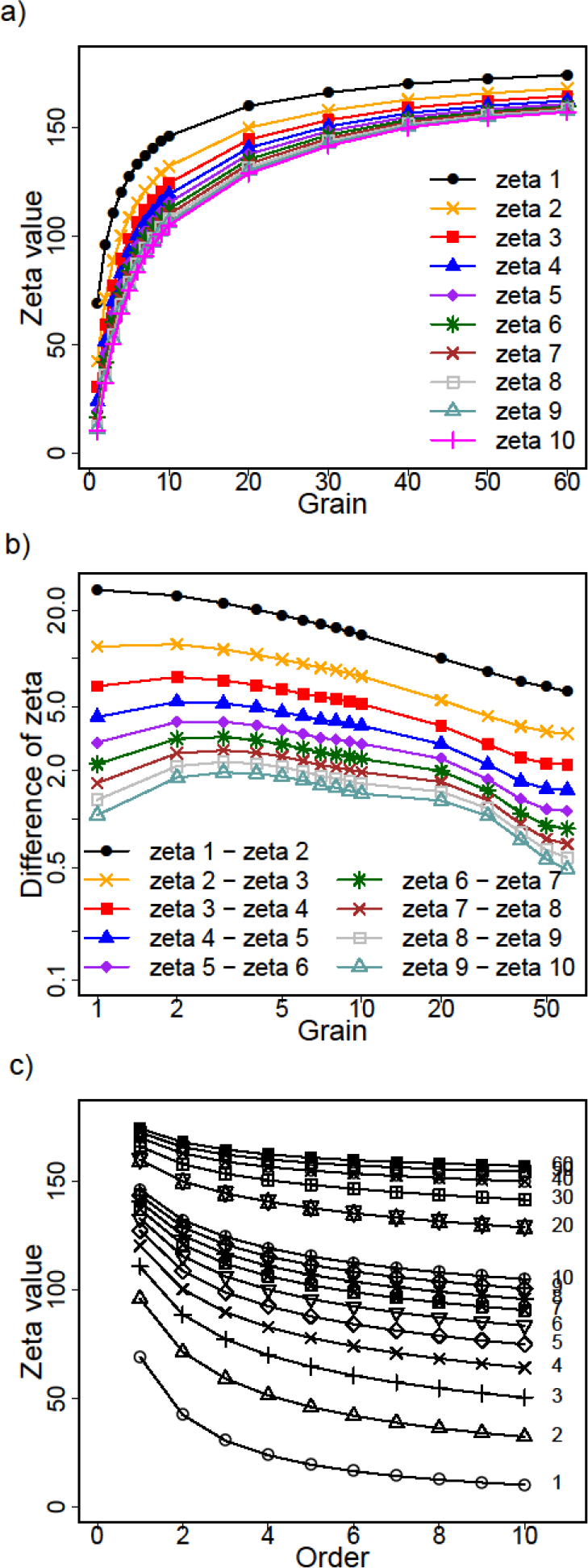
Scale dependence of ζ_1_ to ζ_6_ for the bird data by aggregating 1 to 60 cells based on minimum distance (Figure 6c). a) As the grain increases and cells are aggregated, species share more cells. b) For orders ≥ 2, the difference (ζ_i_ − ζ_i+1_) initially increases with grain, then decreases (Hui et al., 2010). c) The zeta decline from orders 1 to 10 is slightly sharper when aggregating 2 or 3 cells (~1500 km^2^; the grain is indicated on the right) than without aggregating cells (fine grain, corresponding to the ‘1’ zeta decline) or when aggregating more cells (coarse grain, i.e. >4 in this case).

## 7. Concluding remarks

By extending the analyses of compositional change to more than pairwise combinations of assemblages, zeta diversity provides a more detailed understanding of species diversity and a more exhaustive description of community assemblages than using alpha and beta diversity alone. In addition to the clear advantages of obtaining accurate descriptions of biodiversity, such as the possibility to better identify the processes that generates it, zeta diversity also enables the differentiation of the role of common species from rare ones in structuring biodiversity patterns. As we have shown in the examples above illustrating the four different types of analyses currently applicable to zeta diversity applied to bird communities over South-East Australia, considering multiple orders of zeta diversity sheds light on differences in the characteristics and drivers of spatial distribution of common and rare species. It also shows the impact of the spatial resolution at which communities are defined for distinguishing between common and rare species.

The package is also under constant development, and future versions of *zetadiv* will pay special attention to spatially mapping zeta diversity and the parametric form of zeta decline. With increasing recognition of the importance of temporal changes in compositional change (Magurran, 2011) as a consequence of climate change and biotic homogenization (Dornelas et al., 2014), specific functions for temporal decay will be implemented in the future. Current functions can nonetheless already be used to perform such analyses on zeta diversity (e.g. using `Zeta.decline.mc` using the closest assemblages along a temporal gradient). Given the importance of accounting for phylogenetic and functional traits information for the management and conservation of ecological communities (Devictor et al., 2010), phylogenetic and functional measures of zeta diversity will be developed, reflecting similar recent developments for beta diversity (Graham and Fine, 2008; Loiseau et al., 2017). Finally, measures of zeta diversity will be developed for measuring turnover in species interactions.

## Software availability

Name of Software: zetadiv (version 1.1.1).

Year of First Release: 2015.

Developers: G. Latombe, Melodie A. McGeoch, David A. Nipperess, Cang Hui

Maintainer: G. Latombe

Available from the CRAN: https://CRAN.R-project.org/package=zetadiv

## Acknowledgements

We thank Rachel Leihy and Grant Duffy for providing feedback on the functions of the package. We also thank the attendees of the 2015 Ecological Society of Australia and Eco-Stats conferences for fruitful discussions on the concept of zeta diversity. We acknowledge BirdLife Australia as the source of the bird data used here. This research was supported by an Australian Research Council Discovery Project Grant (DP150103017) to MM and CH, and the National Research Foundation of South Africa (grant nos. 81825 and 76912 to CH).

## Supporting information

Appendix A: Supporting tables and figures

Appendix B: Code for reproducibility of examples

